# Novel metrics to measure coverage in whole exome sequencing datasets reveal local and global non-uniformity

**DOI:** 10.1101/051888

**Authors:** Qingyu Wang, Cooduvalli S. Shashikant, Matthew Jensen, Naomi S. Altman, Santhosh Girirajan

**Affiliations:** Bioinformatics and Genomics Program, The Huck Institutes of the Life Sciences; Department of Statistics; Department of Animal Science; Department of Biochemistry and Molecular Biology; Department of Anthropology, The Pennsylvania State University, University Park, PA 16802

## Abstract

Whole Exome Sequencing (WES) is a powerful clinical diagnostic tool for discovering the genetic basis of many diseases. A major shortcoming of WES is uneven coverage of sequence reads over the exome targets contributing to many low coverage regions, which hinders accurate variant calling. In this study, we devised two novel metrics, Cohort Coverage Sparseness (CCS) and Unevenness (U_E_) Scores for a detailed assessment of the distribution of coverage of sequence reads. Employing these metrics we revealed non-uniformity of coverage and low coverage regions in the WES data generated by three different platforms. This non-uniformity of coverage is both local (coverage of a given exon across different platforms) and global (coverage of all exons across the genome in the given platform). The low coverage regions encompassing functionally important genes were often associated with high GC content, repeat elements and segmental duplications. While a majority of the problems associated with WES are due to the limitations of the capture methods, further refinements in WES technologies have the potential to enhance its clinical applications.

## INTRODUCTION

Whole Exome Sequencing (WES) is a high throughput genomic technology that sequences coding regions of the genome selectively captured by target enrichment strategies^1–3^. Target enrichment is achieved by oligonucleotide probes that selectively hybridize and capture the entire coding region of the genome, referred to as the exome^1,3,4^. Since the exome represents approximately two percent of the genome, WES technology provides high coverage at a lower cost and in a shorter time than Whole Genome Sequencing (WGS) technology^5^. From its first successful application in discovering the candidate gene associated with Miller syndrome^6^, WES has been used to study a number of Mendelian^7^ and complex disorders^8–11^. WES is used in the 1000 Genomes Project, the Exome Aggregation Consortium (ExAC), and the NHLBI GO exome sequencing projects to catalog variants in the population and to identify rare variants associated with diseases^12–16^. Since 2011, WES has also been routinely offered as a diagnostic tool in clinical genetics laboratories^17–19^. A recent study reported that in a large cohort of patients referred by a physician to receive genetic testing, 25% of patients received a genetic diagnosis, including diseases such as neurodevelopmental disorders, cancer, cardiovascular disease, and immune-related diseases^20^.

Several target enrichment strategies to capture exomes are available, including the widely used Agilent SureSelect Human All Exon capture kit, Roche NimbleGen SeqCap EZ Exome capture system, and the Illumina TruSeq Exome Enrichment kit. While the basic sample preparation protocols are similar among these platforms, major differences lie in the design of the oligonucleotide probes, including selection of target genomic regions, sequence features and lengths of probes, and the exome capture mechanisms^21–24^. This may lead to some differences in genes captured on each chromosome by the different platforms. In spite of its extensive use, the analysis of WES data still presents considerable challenges. There are significant concerns regarding unevenness of sequence read coverage, which affects downstream analysis. For example, even in samples with high average read depth (>75X), some regions are captured poorly (with coverage as low as 10X), potentially resulting in missed variant calls^25^. Similar issues with uneven coverage can also affect studies with target sequencing strategies, where genomic regions with low read coverage (5X) has decreased sensitivity for detecting variants than regions with higher coverage (20X)^3,4,26^. Studies examining the overall quality of WES data have focused on comparing the performance of a single DNA sample or a small number (n≤6) of samples in different capture technologies^22,23,27^. While these studies have focused on the GC content and overall coverage differences between different platforms, the intra-platform variation in sequence coverage, characteristics of the low-coverage regions, and variation of coverage across the exome have not been quantitatively evaluated. We undertook a comprehensive assessment of sequence coverage in the human exome, and examined the variance in read depth both between samples and across the exome. We evaluated the sequence content and characteristics of the genomic regions contributing to systematic biases in exome sequencing using WES data from a total of 169 individuals obtained from three different platforms. Our study provides quantitative metrics for systematic analysis of different parameters that could potentially impact WES analysis, and confirms the association between low coverage regions and occurrence of duplicated sequences and high GC content.

## RESULTS

To assess the coverage distribution of reads, we selected 169 out of 184 exome sequence samples obtained from NimbleGen, Agilent and Illumina TruSeq platforms, with an average target coverage of at least 75X (**Figure S1, Table S1**). The coverage at specific positions in the exons varied among different samples for each of the three platforms tested. For example, the average read depth of exon 16 of *TP53BP2* in two samples sequenced with NimbleGen at a similar average coverage (92X) was 48X and 92X, respectively. This inconsistency in coverage distribution resulted in a range of 10X-500X read depth at several regions in the exome when multiple samples were run on the same platform. Such regions of highly variable read coverage mapping within disease-associated genes can affect the accuracy of current variant calling algorithms in genetic studies. To characterize the distribution of sequence reads along the exome, we developed two metrics, Cohort Coverage Sparseness (CCS) and Unevenness (U_E_) scores. The CCS score provides an assessment of coverage of all exons across the genome (global) in the given platform, while the U_E_ score provides an assessment of coverage of a given exon (local) across different platforms.

### Coverage deficiencies determined by CCS Score

Read coverage of a position in the genome is considered deficient if the number of reads mapped to that position is less than 10 reads^21^. The CCS score is defined as the percentage of low coverage (<10X) bases within a given exon in multiple WES samples. The CCS score is estimated by first calculating the read depth for each base position in a given exon, and then determining the median percentage of samples with low coverage at that region (see Methods). The resulting CCS score may vary between 0 and 1, with high CCS scores indicating low sequence coverage. We plotted CCS scores for all genes against the chromosome positions in a modified Manhattan plot (Figure 1). As shown in the figure, a majority of genes (>88%) were clustered in the low CCS score region (<0.2), indicating good coverage in all three platforms (Figure 1A-C). The remaining genes with CCS scores >0.2, which contain low coverage regions, were scattered throughout the plot. In those low coverage genes, CCS score >0.5 indicates nearly half of the regions have less than 10X read depth. The distribution of CCS scores is skewed to the right when plotted as a histogram (Figure S2), consistent with the pattern shown in Figure 1. Further, the data generated from Illumina TruSeq have the lowest percentage of low coverage genes (~7%), compared with data generated by the capture kits from NimbleGen (~10%) and Agilent (~11%), as shown in Table 1. The differences in these percentages could be due to differences in the design of probes that target the exome in these platforms^23^. **Table S2** lists all autosomal low coverage genes identified in the three WES platforms compared to a WGS dataset.

**Figure 1:**
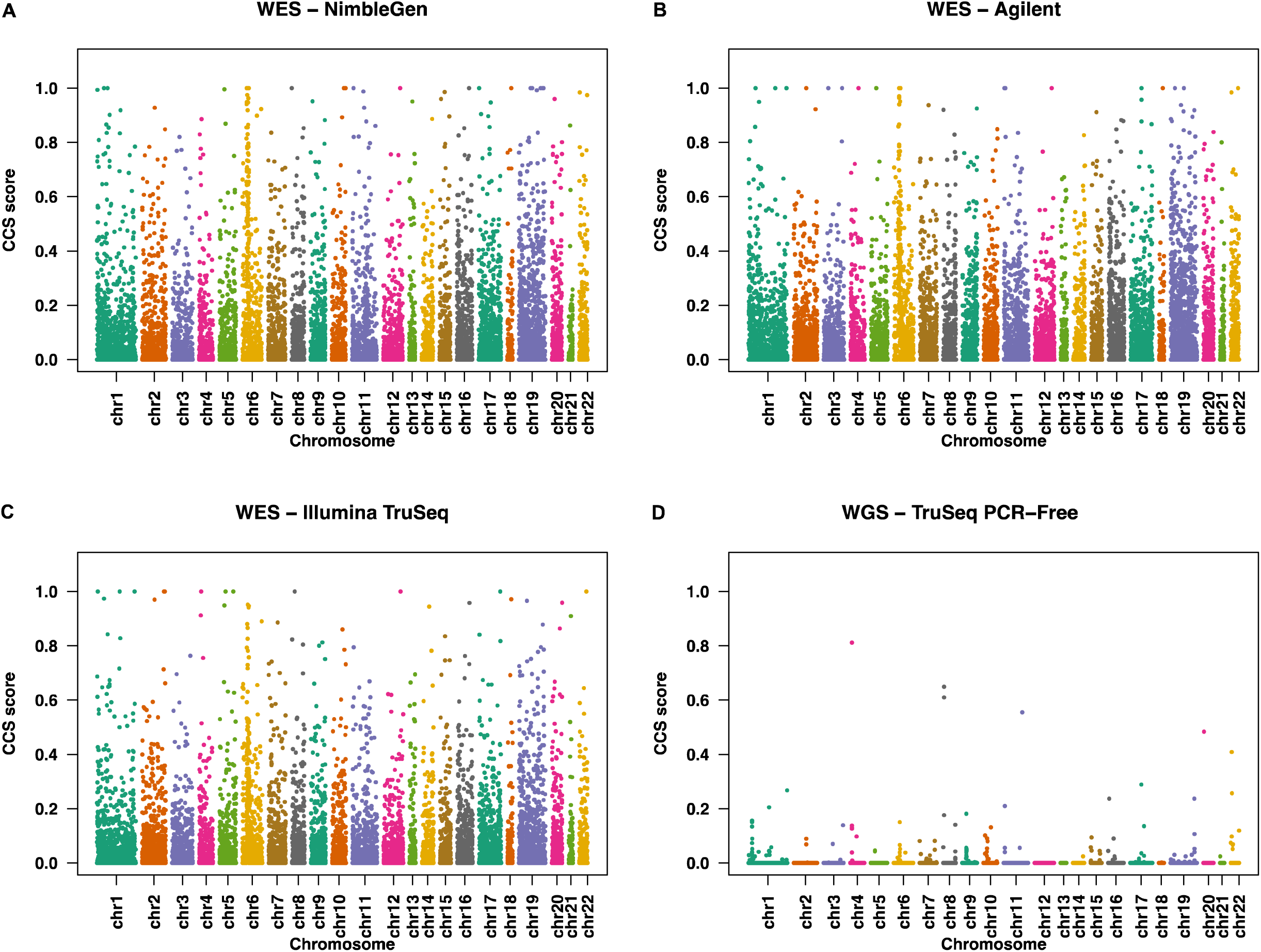
CCS scores of targeted RefSeq genes along the whole chromosome in WES and WGS datasets. The CCS values are plotted along the length of each chromosome in a modified Manhattan Plot for WES datasets obtained from (**A**) NimbleGen, (**B**) Agilent, (**C**) Illumina TruSeq, and (**D**) WGS dataset from 1000 Genomes project.

**Table 1.** Low coverage genes with high CCS scores in three different datasets

| **Platforms** | **Genes\*** | **Genes with CCS > 0.2 (%)** | **Genes with CCS > 0.5 (%)** |
| --- | --- | --- | --- |
| **NimbleGen** | 18,024 | 1,819 (10%) | 428 (2%) |
| **Agilent** | 17,780 | 2,025 (11%) | 374 (2%) |
| **Illumina TruSeq** | 17,866 | 1,252 (7%) | 228 (1.3%) |

\*Gene sets in different platforms were defined as described in the Methods section.

The low coverage regions (CCS>0.2) varied significantly across all chromosomes in the three platforms analyzed (**Table S3**; chi-square test p-value <2.2×10^−16^). Chromosomes 6 and 19 have a higher proportion of low coverage genes compared to other chromosomes. A low coverage gene cluster on chromosome 6 (cytobands 6p21.33 and 6p21.32) corresponded to the genes encoding human leukocyte antigen, which are known to be polymorphic and have alleles showing high sequence identity^28,29^. Similarly, chromosome 19 is known to carry a high proportion of tandem gene families, repeat sequences, and segmental duplications (SD)^30^. These sequence features potentially affect accurate mapping of reads, leading to low coverage regions.

### Coverage non-uniformity determined by U_E_ Score

The U_E_ score provides a measure of non-uniformity calculated after smoothing the coverage distribution curves and identifying peaks and troughs along the curve (Figure 2, see Methods). It is calculated based on the number and structural features (height, width, base) of the coverage peaks. The U_E_ score increases with an increase in the number and relative height of peaks within a given exon. For regions with uniformly distributed coverage, the U_E_ score is 1; for regions with uneven coverage, the U_E_ score is greater than 1. To illustrate how U_E_ score varies among different platforms, we chose the last coding exon of *ZNF484*, which had high (CCS scores < 0.01) but variable (U_E_ score 72.2-141.9) coverage across the different platforms, indicating inconsistent coverage (Figure 3).

**Figure 2:**
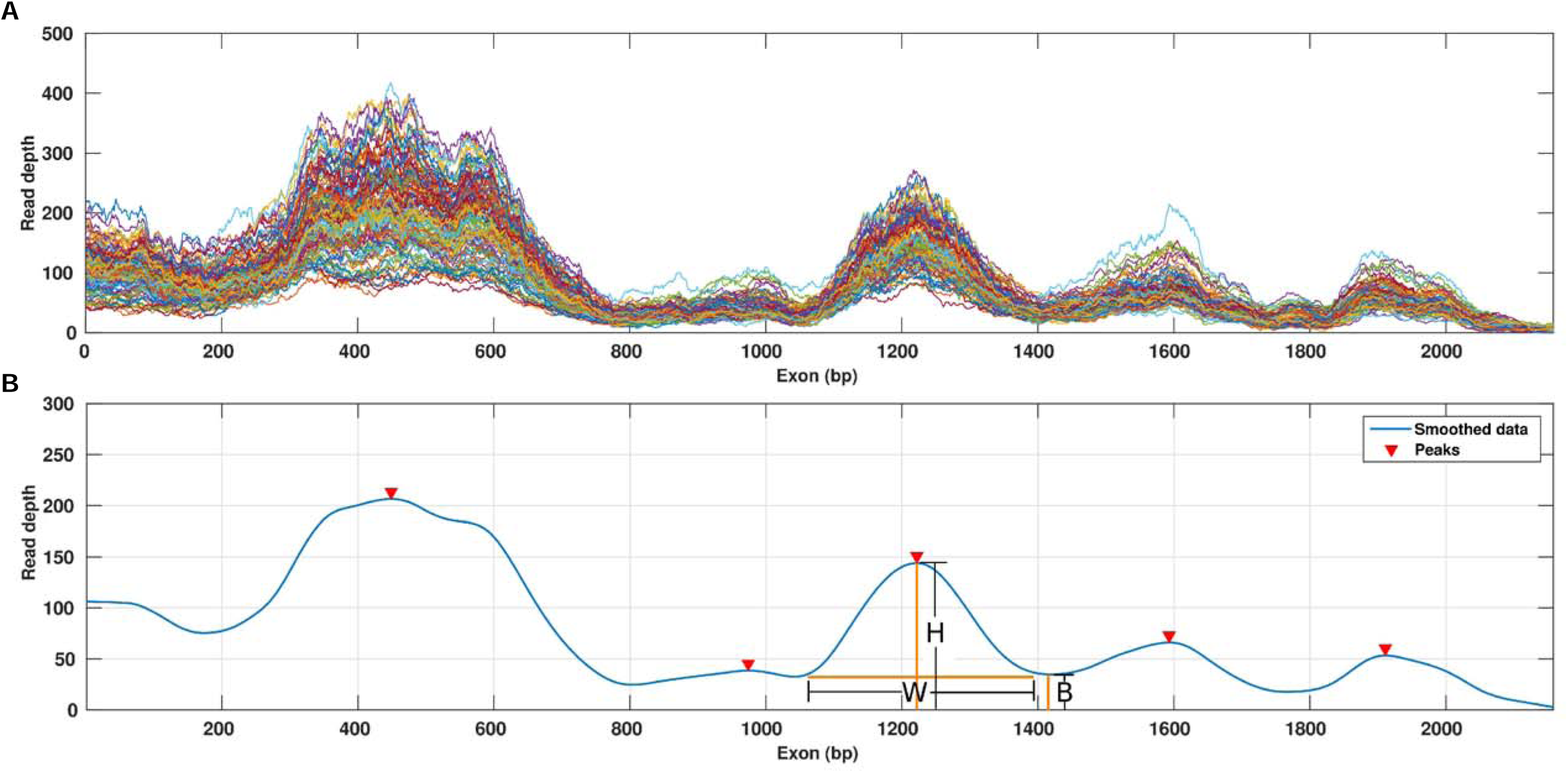
Characterizing the coverage distribution with the Unevenness (U_E_) score. (**A**) The coverage distribution from multiple samples is plotted against the exon length. (**B**) The smoothed median coverage plotted against the exon length, obtained by first calculating median coverage for each position and then using LOWESS smoothing. Peaks and troughs were then identified by using a local optimization algorithm. Arrows indicate peaks identified in the curve: B, base, W, width and H, height of the peak, L^R^, length of the region analyzed

**Figure 3:**
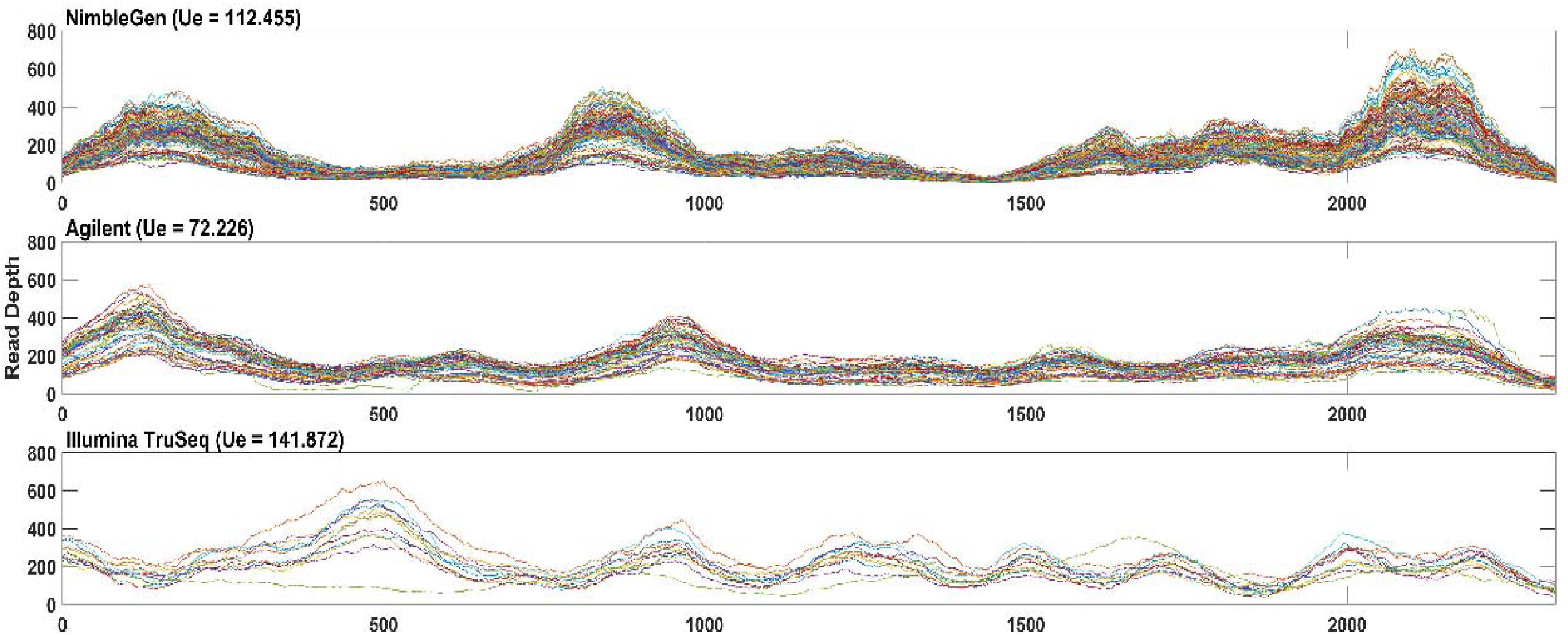
Base coverage distribution along the length of the last coding exon of gene *ZNF484* from WES datasets obtained from (**A**) NimbleGen, (**B**) Agilent, and (**C**) Illumina TruSeq.

We also observed a positive association between exon length and the U_E_ score in all three platforms as shown in the scatter plot (Figure 4). The Pearson correlation coefficient for all platforms was ≥0.7 (NimbleGen, 0.80, Agilent, 0.71; TruSeq, 0.70). The U_E_ score was significantly different for longer exons (>400 bp) among the three platforms tested (Friedman test, p-value <2.2×10^−16^). When the coverage distribution of the neighboring exons was examined, we found inconsistencies in the rank order of read coverage among samples tested on the same platform (**Figure S3**). Since most of the current methods for calling copy number variations (CNVs) are based on detecting continuous depletion or enrichment after normalizing for coverage of adjacent exons^31–33^, this lack of consistency of rank orders between closely occurring exons could affect CNV calling from exon read depths.

**Figure 4:**
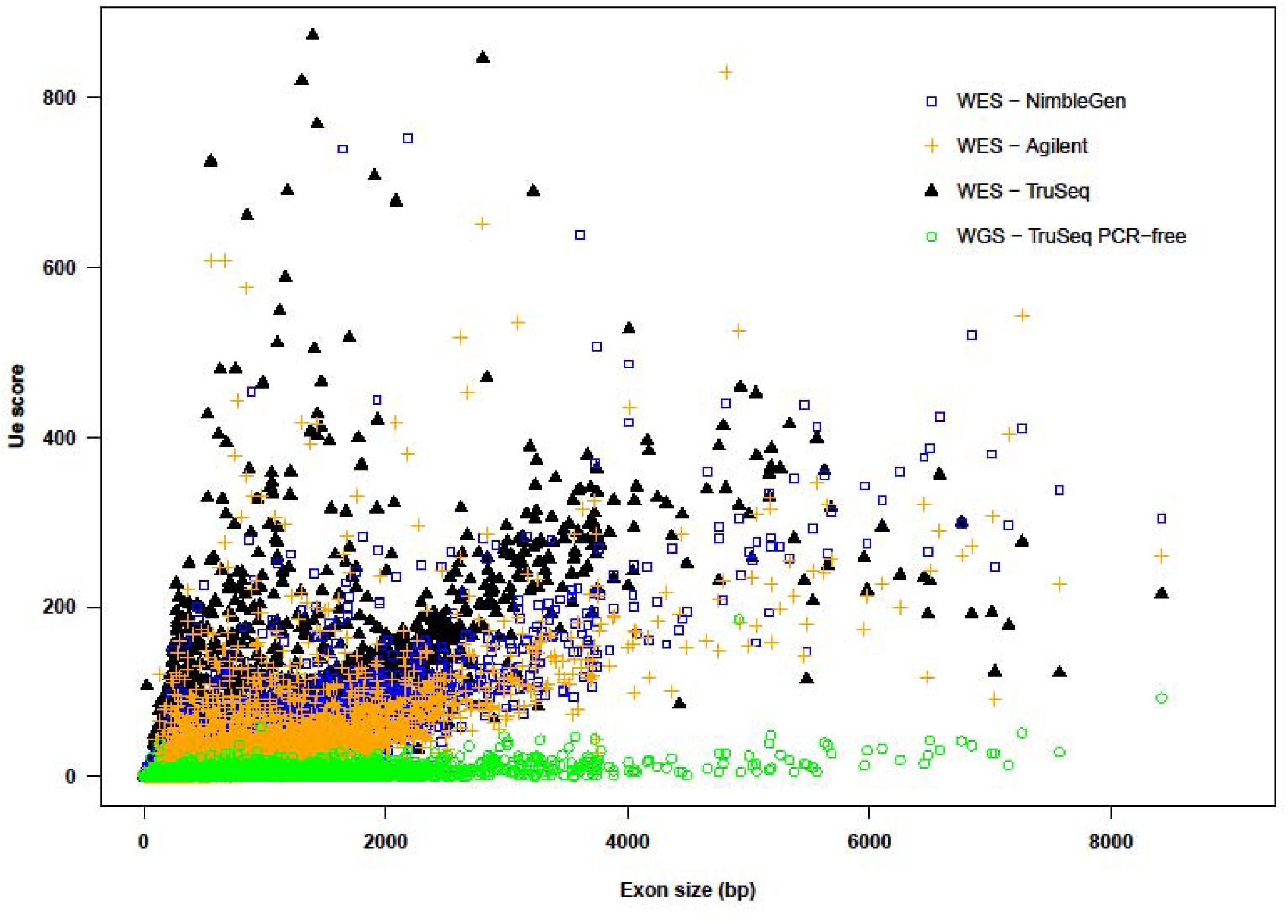
Scatterplot of Unevenness (U_E_) scores against exon size in WES and WGS datasets.

### Low coverage regions are enriched within repeat elements

Since chromosomes 6 and 19 contain repeat elements and clustered gene families that accounted for specific regions showing low coverage, we checked if these sequence features globally correspond with the occurrence of low coverage. In one study, SD regions from the Agilent WES dataset and the Segmental Duplication Database (http://humanparalogy.gs.washington.edu) were cross-referenced with the low coverage genes obtained from our analysis, and examined for the percentage of reads that mapped unambiguously to off-target regions. The percentage of off-target reads with multiple hits in SD regions (50%) was about double that of those found in unique regions (26%). These results suggest that the mapping method may also contribute to the missing coverage.

Because SD regions share similar features with repeat elements, we next tested if low coverage regions were associated with underlying repeat sequences. We first examined the 358 bp exon 1 of *MAST4*, which contains a 22 bp low complexity repeat element predicted by RepeatMasker (http://www.repeatmasker.org)^34^, in four samples sequenced with the NimbleGen platform with highly variable overall coverage (75X-200X). As shown in Figure 5, the coverage was high (30X) for sequence base positions 1-200 in samples with high average coverage. The read depth at each nucleotide in this region remains high. In contrast, the coverage falls dramatically (to <10X) between base positions 200-358, with corresponding low read depth at each nucleotide in this region. The low coverage region is greater than a typical exome capture probe size (50-150 bp), indicating that influences of a repeat element can extend beyond its location. This shows that even when using a platform with high variability in coverage across samples, some genomic regions associated with repeat sequences can consistently show low coverage.

**Figure 5:**
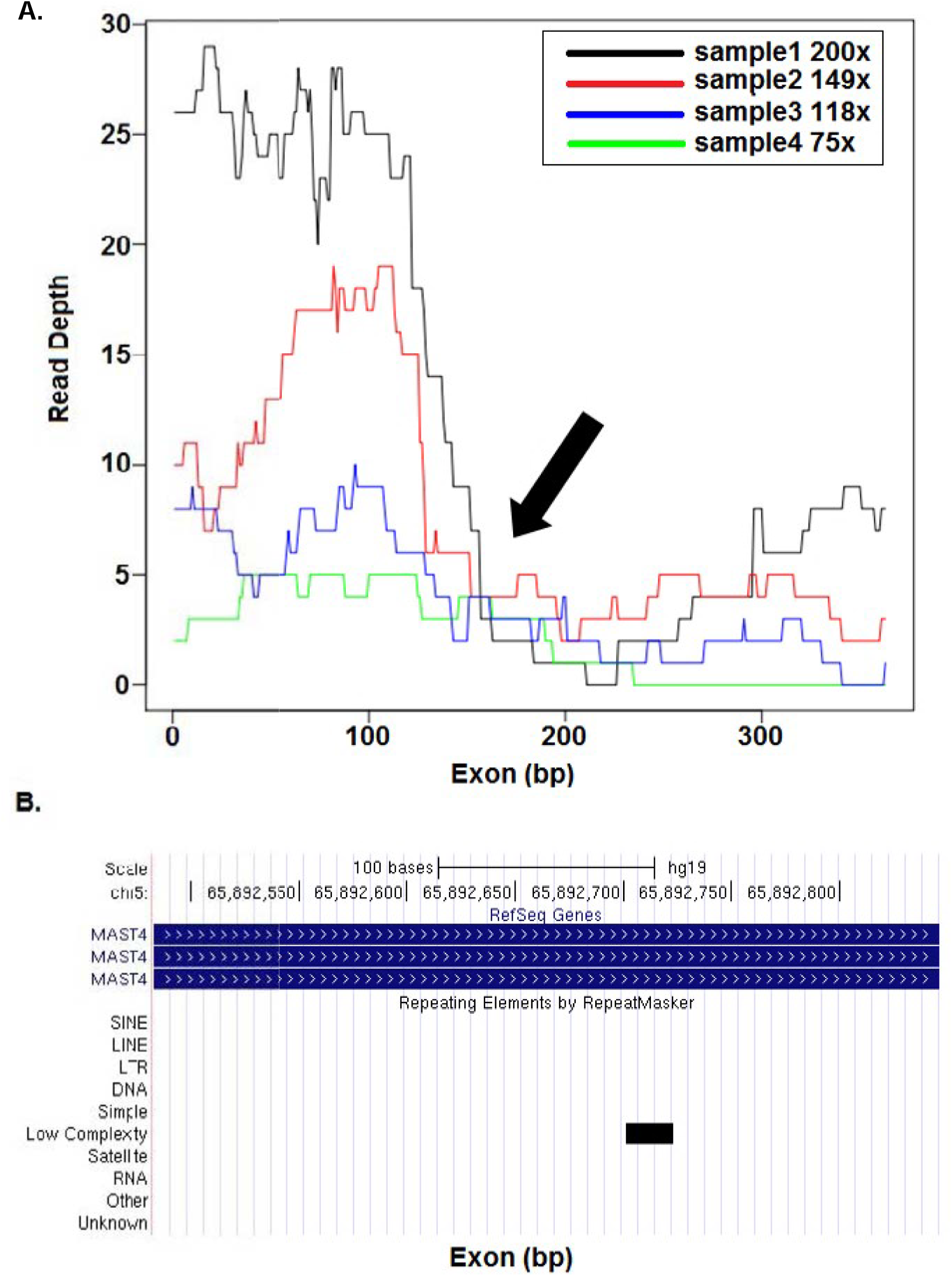
Concurrence of repeat elements and coverage sparseness. (**A**) Base coverage distribution along the length of the first coding exon of *MST4*. WES samples from the NimbleGen platform with different average coverage ranging from 75X to 200X are shown in different colors. Arrow indicates the point at which coverage falls sharply. (**B**) UCSC browser screen shot of MST4 genomic region, black bar indicates the position of the repeat element.

To test for a global association between the occurrence of repeat elements and low coverage, we examined all the troughs (from U_E_ score calculations) in coverage in the 169 samples studied. Repeat elements predicted by RepeatMasker coincided with extremely low troughs (median coverage <10X) for a majority of the exons (60% to 69% of exons tested on the three platforms), especially for exons >200 bp in size. For example, as shown in **Figure S4**, multiple repeat elements occur in the extremely low trough region (positions 1 to 400 bp) in the last coding exon of *CASZ1.* This suggests that coverage troughs are highly associated with repeat sequences. However, we also find that several troughs contain repeat elements that are not annotated by RepeatMasker when default parameters are used (**Figure S5**).

### Negative association between GC content and sequence coverage

Since GC content is one of the major factors contributing to low coverage in WES data ^23,35–37^ we investigated whether there is an association between GC content and CCS score. We used a density curve plot and a modified density plot, which visualizes the distribution of GC content of genes with the corresponding CCS scores (Figure 6, **Figure S6**). Based on GC content, we were able to clearly distinguish exons with CCS score <0.2 from exons with CCS score >0.2. Thus, high GC content correlated with high CCS scores, and therefore with lower coverage regions. Exons with good sequence coverage (CCS<0.2) were clustered in the intermediate GC content regions (between 30-70%). In contrast, a higher density of low coverage exons (CCS>0.2) was observed in regions with relatively high GC content (>70%). Very few good coverage exons were present in regions with less than 20% or more than 80% GC content (**Table S4**). However, some poorly covered exons (CCS>0.8) were found in the intermediate GC content regions (<70% and > 30%), suggesting that there may be other factors contributing to the low coverage within these regions.

**Figure 6:**
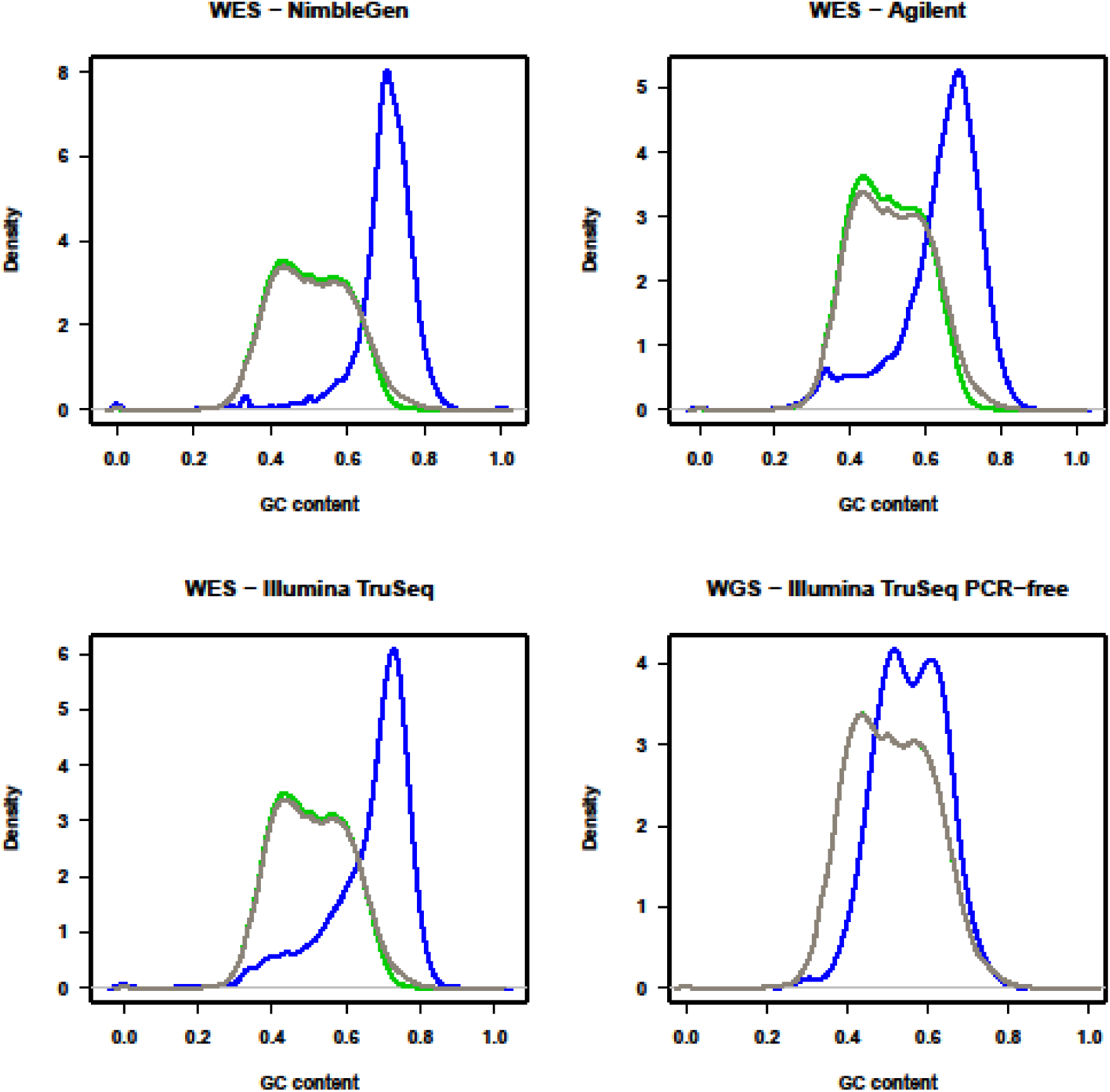
The probability density curves showing GC content in sets of genes with different coverage. The distribution of GC content of all genes (black), high coverage genes with CCS score < 0.2 (green), low coverage genes with CCS score > 0.2 (blue) are represented.

### Low coverage regions contain functionally relevant genes

To examine whether low coverage regions had functional significance, we conducted Functional Disease Ontology Annotations (FunDO)^38^ of 832 low coverage genes that were common to all three platforms (Figure 7A). Result showed enrichment of genes implicated in leukemia, psoriasis, and heart failure (Figure 7B). We also examined the coverage of genes that American College of Medical Genetics and Genomics (ACMG) recommends for pathogenic variant discovery and clinical reporting^39,40^. Of the 59 genes examined, six genes, including, *KCNH2, KCNQ1, SDHD, TNNI3, VHL*, and *WT1*, mapped within low coverage regions in one or more samples (with average coverage >75X) (**Figure S7**). These results suggest that low coverage regions within functionally important genes could affect variant discovery and subsequent clinical diagnosis.

**Figure 7:**
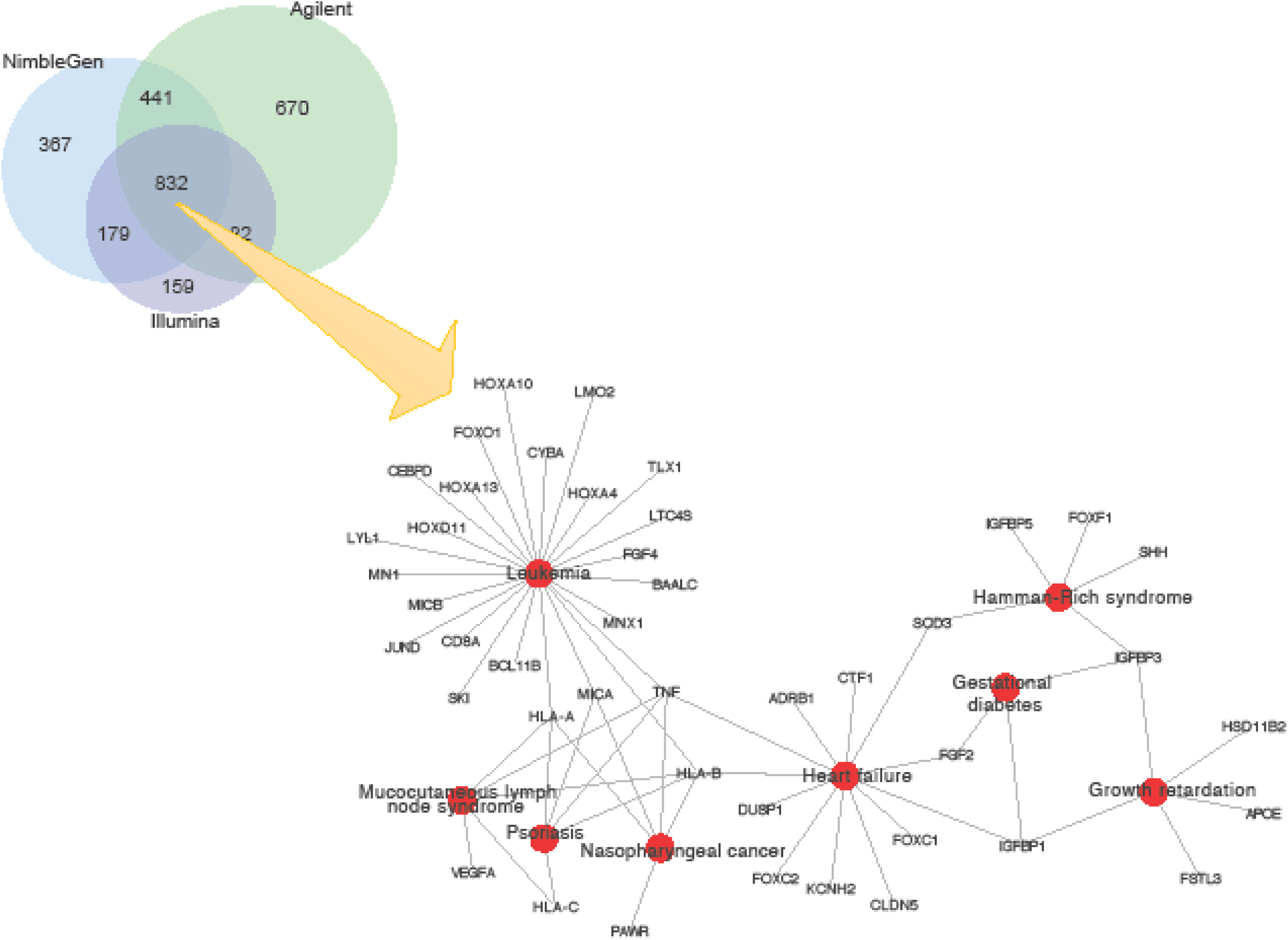
Genes with low coverage in three different datasets. (**A**) Venn diagram showing number of low coverage genes (CCS score > 0.2) across three different platforms. There are 832 genes with low coverage in common across all platforms. (**B**) Network diagram showing disease ontology analysis of the 832 low-coverage genes showing associations with leukemia, psoriasis, heart failure, and mucocutaneous lymph node syndrome.

### Low coverage is more of an issue for WES than WGS based platforms

We examined differences in the exon coverage between the WES and WGS datasets to test whether mapping issues were common to both platforms. We examined the CCS scores for all genes (Figure 1D), and the U_E_ scores (Figure 4) and GC content (Figure 6D) for all exons. For WGS datasets with an average coverage of about 60X (**Table S2**), only 15 genes with high CCS scores were observed in the modified Manhattan plot (Figure 1D). In contrast, for WES platforms with an average coverage of >75X, over 1,000 genes with high CCS scores were observed (Figure 1A-C). Similarly, the U_E_ score for all exons in the WGS analysis was significantly lower compared to the WES analysis (Friedman’s test, p<2.2×10^−16^). As shown in Figure 4, the U_E_ score increased slightly with increased size of the exon. To examine how GC content affects coverage in the WGS dataset, we generated probability density curves for GC content for different gene groups (Figure 6). In all the three WES platforms, we found a dramatic shift in the density plots for high coverage genes compared to low coverage genes based on GC content. In comparison, for the WGS platform, the shift in GC content between the exons with high and low CCS scores was minimal. Thus, GC content appears to have less influence on sequence coverage in WGS than WES analysis. These results suggest that problems of low coverage are specific to WES platforms, and these limitations could be contributed by technological differences (such as capture bias) in WES ^41–43^. Therefore, the methods developed for WGS analysis require further modifications before application to WES platforms.

## DISCUSSION

WES is a powerful clinical diagnostic tool for identifying disease-associated variants in patients^44,45^. Since most known Mendelian disorders are associated with mutations in the coding regions, focusing on sequencing exomes rather than whole genomes is efficient in terms of time, expense and coverage^5^. Recent studies have successfully used WES technology to identify variants that strongly correlate with disease phenotypes^19,20,46,47^. However, high-resolution examination of different WES datasets shows uneven coverage along the length of exons, which could cause possible problems in variant calling analysis. This affects identification of de novo variations that may be clinically important. In this study, we systematically examined different parameters that could potentially impact WES analysis and identified key issues associated with sequence architecture contributing to the low coverage.

We analyzed WES data captured by three major platforms: NimbleGen CapSeq V2, Agilent SureSelect V2 and Illumina TruSeq. All three platforms are based on similar target enrichment protocols and cover >95% of RefSeq^48^ coding region with >88% low CCS scores. These platforms differ from one another in the layout and length of probes. Most importantly, while NimbleGen uses overlapping probes, Agilent uses tiling probes, and Illumina uses gapped^21–24,37^. This difference in probe design contributes towards inconsistency in the coverage, creating systematic biases and preventing the combining of datasets from different platforms for SNV and CNV detection^21–24,37^. This is a major concern, especially with large cohort studies involving multiple centers using different platforms. While the heterogeneity of the cohorts and the differences in the number of samples in each cohort could potentially affect our conclusions on the evaluation of the WES platforms (for example, cancer samples in the TruSeq data versus samples from the developmental disorder cohort in the NimbleGen data), we find a common set of genes that are affected by low coverage irrespective of differences in protocols, tissue samples, and sequencing and platform biases. In fact, 832 genes are covered at low depth in all the three platforms (Figure 7).

Even in the data generated by a single platform, the coverage distribution varied among different exons. Several factors including size of the exon, GC content, presence or absence of repeat elements, and segmental duplications affect the coverage. The uniformity of coverage distribution decreases for longer exons. For a given exon, the pattern of coverage varied among different platforms even for genes with high coverage (Figure 3), making it difficult to normalize the background coverage for CNV calling. As shown in our study, coverage in regions with extremes of GC content (<30%; >70%) was low reflecting poor capture efficiency. In contrast, no such correlation was observed for regions with moderate GC content (30-70%). The coverage was also affected by the presence of repeat elements. Even simple repeat elements as small as 22 bp may contribute to low coverage. Those platforms, which use RepeatMasker in probe design, are likely to miss exonic regions containing repetitive elements. We also note that while the CCS and U_E_ metrics are useful in identifying areas of the genome with uneven coverage across multiple samples, they will miss genomic regions that have consistently low coverage across all tested samples.

Several studies comparing technologies have found WGS to be superior to WES^41,49,50^. The parameters affecting coverage including GC content, uniformity of the coverage, and read depths were examined. Our results are consistent with these findings. Although WGS data has lower average coverage, the coverage depth along exons is more uniformly distributed. WGS data have fewer sparse regions, which may contribute to lower numbers of false negative variant calls. However, with more than 100,000 exomes sequenced to date, WES has become the major genetic tool in several diagnostic centers^45,51^. A thorough understanding of the limitations of each of the WES platforms is thus important. Modifications in the design of the targeted sequence capture technology and improvements in mapping algorithms are essential for accurate calling of variants and filling the gaps in heritability estimates of genetic disease.

## MATERIALS AND METHODS

### Datasets and Pre-processing

#### Datasets

WES data generated from NimbleGen SeqCap, Agilent SureSelect, and Illumina TruSeq were obtained for analysis from the dbGaP (the database of Genotypes and Phenotype)^52^ and SRA (Sequence Read Archive) databases^53^. WGS sample data were obtained from the 1000 Genomes Project, Phase 3 analysis ^54^ Detailed information for each dataset is listed in **Table S1**.

#### Mapping of reads

All SRA files of exome samples downloaded from different databases were first converted to FASTQ files using the SRA Toolkit^53^. Then, raw sequence reads were mapped to the reference genome using Bowtie2 version 2.1.0, with default parameters^55^. We used the hg19/GRCh37 assembly of the human genome as the reference sequence throughout the analysis, and SAMtools version 1.2 was used to sort reads and remove PCR duplicates^56^. The WGS dataset was downloaded as raw FASTQ files from the 1000 Genomes servers, and were mapped using Bowtie2 version 2.1.0, with parameter −X 1500 to adjust the maximum insert size for valid paired-end alignments^55^. Mapping statistics for each data set, including average library size, number of mapped reads, percent PCR duplicates, and coverage are listed in **Table S5**.

Reads with low mapping qualities were retained in order to identify all regions with low coverage. We note that Bowtie2, like other alignment software, randomly assigns reads to a location if multiple optimal locations are identified^55^. As we retained such reads, coverage of highly homologous genomic regions may differ between multiple alignments of the same sample. However, when we compared the average mappability of each region^57^ to both CCS and U_E_ metrics, we found no statistical correlation to our metrics (**Figure S8**).

#### Exome and gene sets annotation

Overlapping exons of the same gene from different transcripts were merged to create a consensus exome annotation file from the RefSeq database using an inhouse pipeline. Target regions obtained from three different capture kits were mapped and intersected with the consensus exome annotation file. These target regions were then sorted by genomic coordinates on chromosomes and defined as target exon regions. For WGS data analysis, all exons in the consensus exome annotation file were used as target exon regions. Only autosomal chromosomes 1 to 22 were included in this analysis.

#### Coverage calculation

BEDTools version 1.v2.18.2^58^ was used to calculate the single base pair coverage of all BAM files at all positions covered in the exome annotation file.

### Metrics to Evaluate Low Coverage Regions

#### CCS score

The CCS score for all exons and genes was calculated with the ExomeCQA program. The CCS score provides a measure of percentage of base pairs in a given region for which coverage is less than 10 reads in multiple samples (see **Figure S9**). The CCS score is calculated by the formula:

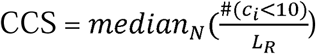

where, *N* is the total number of samples; *#* is the count of the genomic positions with read depth less than 10X; *c_i_* is the coverage at genomic position *I;* and *L_R_* is the length of the region of interest.

#### U_E_ of coverage

The U_E_ metric was introduced to measure the non-uniformity of the coverage over targeted regions, and is calculated using ExomeCQA. The score is based on peaks and troughs in the coverage. For a targeted region, the median coverage at each genomic position is computed from all the samples. The median coverage is smoothed against base position using the LOWESS (locally weighted scatterplot smoothing)^59^ method with an empirically selected span 0.03, which dampens the locus-to-locus variability of the curve so that local trends can be detected. Here we defined “span” as a parameter that represents the proportion of the total number of base positions of the exon region that contribute to each locally fitted coverage value. We used a percentage-based span parameter instead of a constant base-pair parameter in the LOWESS method, as we wanted to consider the length of the peak in relation to the length of the exon after smoothing. Using a percentage-based span will allow smaller peaks and troughs in the shorter exon, while in longer exons small peaks and troughs will be smoothed over. This justifies considering all peaks when calculating the U_E_ formula. Peaks and troughs of the median coverage were identified by a Hill Climbing local optimization algorithm^60^, which was implemented by scanning all base positions and identifying the positions without increasing or decreasing neighbors in the smoothed coverage curve. The height and width of the peaks were then used to calculate the unevenness score by the following formula:

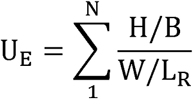

where U_E_ is unevenness score, N is number of peaks, H is height of the peak, B is base of the peak, and W is width of the peak.

We note that the library size is not considered in U_E_ score calculation. When the local coverage is proportional to the library size, samples with (for example) double the coverage should have peaks that are twice as high. In the high coverage regions, we expect the coverage to be proportional to the library size. If the low coverage regions are also proportional to the library size, then height and base will change proportionally and U_E_ will remain the same. However, we have already seen (Figure 6) that there are some exonic regions in which no reads map. In that case, the height of the peak increases but the height of the base does not, increasing U_E_. This adaptation to library size seems appropriate, since in the former case the relative size of the peaks is maintained, whereas in the latter case, the higher peaks do actually lead to less even coverage.

#### Program ExomeCQA

ExomeCQA was written to calculate different metrics of cohort exome sequencing data from coverage files input. This program was written in C++ and can be downloaded at http://exomecqa.sourceforge.net.

### Statistical Methods

All the statistical tests were conducted in R. The Manhattan plots of CCS scores in all chromosomes were generated with the “GWASTool”^61^ package. The density plot of GC content and low coverage regions was generated with the package “GenePlotter”^62^. Both packages are hosted in Bioconductor^63^.

## Acknowledgements

Acknowledgements: We thank Debmalya Nandy and Emily Huber for critical reading of the manuscript. This work was supported by a Basil O’Connor Award from the March of Dimes Foundation (#5-FY14-66), R01-MH107431, a NARSAD Young Investigator Grant from the Brain and Behavior Research Foundation, and resources from the Huck Institutes of the Life Sciences (SG), T32-GM102057 and The Pennsylvania State University Experiment Station grant AES 4586 (CCS).

We acknowledge the NIH data repository, the contributing investigators, and the associated primary funding organizations for providing access to the datasets used in this study. We thank NIH Genomic Study Datasets, Christopher A. Walsh, MD, PhD. Children's Hospital Boston, Boston, MA, USA (PI of the dataset), and their funding resource: U54 HG003067. National Human Genome Research Institute, National Institutes of Health, Bethesda, MD, USA; the Department of Health Sciences, University of Milano-Bicocca, Monza, Italy (for SRP028277); Liver Cancer Institute, Zhongshan Hospital, Fudan University, Grant: ID 81272725, "Identification of driving mutations and singling pathway alterations in intrahepatic cholangiocarcinoma", National Natural Science Foundation of China (for SRP025150); McGill University, Cancer Research Society, Hungarian Scientific Research Fund (OTKA) contract T04639, Canadian Institutes of Health Research grant 102684, and National Research and Development Fund (NKFP) (for SRP032767). Funding support for the dataset in “Sporadic autism exomes reveal a highly interconnected protein network of de novo mutations” (phs000482.v1.p1) was provided by the Simons Foundation Autism Research Initiative (SFARI 137578 & 191889), NIH HD065285 and the Howard Hughes Medical Institute. Data was originally reported by O'Roak et al. 2012, PMCID: PMC3350576. We are grateful to all of the families at the participating SFARI Simplex Collection (SSC) sites, as well as the principal investigators (A. Beaudet, R. Bernier, J. Constantino, E. Cook, E. Fombonne, D. Geschwind, E. Hanson, D. Grice, A. Klin, R. Kochel, D. Ledbetter, C. Lord, C. Martin, D. Martin, R. Maxim, J. Miles, O. Ousley, K. Pelphrey, B. Peterson, J. Piggot, C. Saulnier, M. State, W. Stone, J. Sutcliffe, C. Walsh, Z. Warren, E. Wijsman).

## Author Contributions

Q.W. and S.G. conceived the problem. Q.W., S.G., N.S.A., C.S.S. contributed to the design of computational approaches. Q.W. performed all the analysis, developed algorithms and the ExomeCQA program. M.J. performed mapping of WGS data. Q.W., C.S.S., and S.G. wrote the manuscript, and N.S.A., M.J. and S.G. contributed to the revisions.

